# Canary in the cardiac-valve coal mine. Flow velocity and inferred shear during prosthetic valve closure –predictors of blood damage and clotting

**DOI:** 10.1101/2022.06.23.497372

**Authors:** Lawrence N. Scotten, Rolland Siegel, David J. Blundon, Marcus-André Deutsch, Terence R. P. Martin, James W. Dutton, Ebrahim M. Kolahdouz, Boyce E. Griffith

**Author notes:** Correspondence to: Lawrence N. Scotten. Dipl. T., Independent Consultant, Victoria, BC, V8P 4V4, Canada. See appended Supplement S9. In an early coal mining era, small songbirds accompanied workers in subterranean extraction sites. As these avian species are highly sensitive to changes in ambient air quality, they provided early warnings of impending hazardous breathing conditions.

## Abstract

**Objective:** To demonstrate a clear link between predicted blood shear forces during valve closure and thrombogenicity that explains the thrombogenic difference between tissue and mechanical valves and provides a practical metric to develop and refine prosthetic valve designs for reduced thrombogenicity.

**Methods:** Pulsatile and quasi-steady flow systems were used for testing. The time-variation of projected open area (POA) was measured using analog opto-electronics calibrated to projected reference orifice areas. Flow velocity determined over the cardiac cycle equates to instantaneous volumetric flow rate divided by POA. For the closed valve interval, data from quasi-steady back pressure/flow tests was obtained. Performance ranked by derived maximum negative and positive closing flow velocities, evidence potential clinical thrombogenicity via inferred velocity gradients (shear). Clinical, prototype and control valves were tested.

**Results:** Blood shear and clot potential from multiple test datasets guided empirical optimization and comparison of valve designs. Assessment of a 3-D printed prototype valve design (BV3D) purposed for early soft closure demonstrates potential for reduced thrombogenic potential.

**Conclusions:** The relationship between leaflet geometry, flow velocity and predicted shear at valve closure illuminated an important source of prosthetic valve thrombogenicity. With an appreciation for this relationship and based on our experiment generated comparative data, we achieved optimization of valve prototypes with potential for reduced thrombogenicity.

**Competing Interests:** None declared.

**Financial Disclosure:** This research has been done on a pro bono basis by all authors.

**Graphical Abstract:** Visualization of water jetting through closed mechanical heart valve under steady flow. Under pulsatile conditions, similar jet patterns near valve closure and leaflet rebound are likely. Dynamic metrics for several valves assessed in vitro are important in prediction of comparable blood cell damage and potential life-threatening thrombotic outcomes. Red star indicates moment of valve closure.

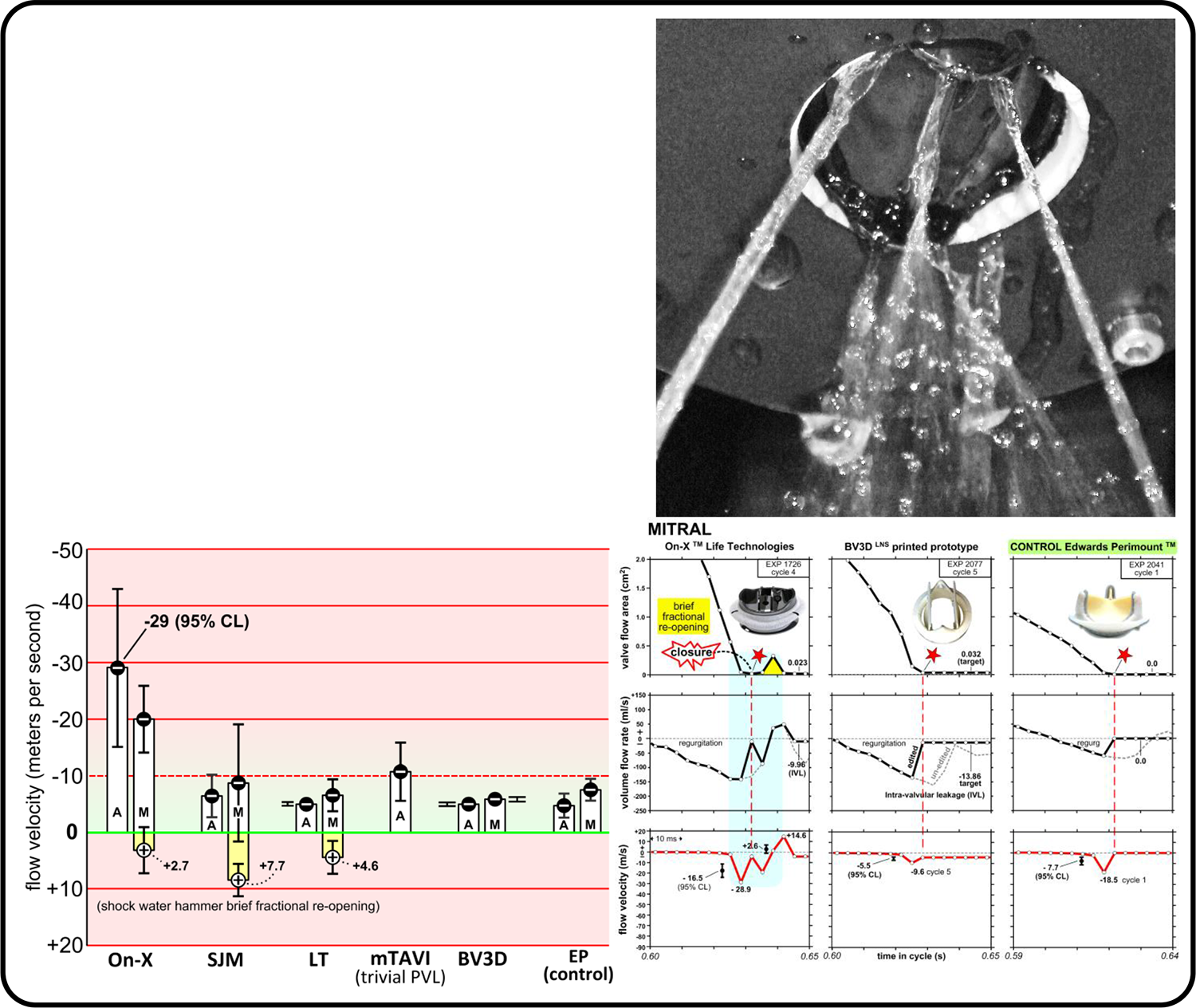

**CENTRAL MESSAGE:** A derived laboratory metric for valve closing flow velocity offers a way to rank valve models for potential blood damage. These results provide new insight and a mechanistic explanation for prior clinical observations where aortic and mitral valve replacements differ in thrombogenic potential and anticoagulation requirement. The study suggests a path forward to design and evaluate novel mechanical valve models for future development. As multiple modifications to mechanical and bioprosthetic valves have not resolved chronic shortcomings related to thrombogenicity and durability, a new development avenue was required to lead to eliminate thrombogenicity in the former and extend durability in the latter.

**PERSPECTIVE:** Prosthetic mechanical valve devices cause blood cell damage. Activation of the coagulation cascade is initiated by dynamic valve function. Design innovation focusing on valve closure behavior may reduce valve thrombogenic potential. **Our study demonstrates that valve design can be empirically optimized with emphasis on that phase.**

**SIGNIFICANCE:** Emphasis on open valve performance has encouraged a long-standing bias while under appreciation of the closing phase vital to identification of potential thrombogenic complications persist. Our multiple data sets are useful in challenging this bias.

Dynamic motion(s) of mechanical valves and derived regional flow velocity are impacted by valve geometry. Focus on valve closure dynamics may lead to the development of potentially less thrombogenic prototype valves. Laboratory experiments support the supposition that valve regional flow velocity is associated with valve thrombogenic potential. This study compares three clinical valves and two experimental prototypes.

## INTRODUCTION AND BACKGROUND

Our interest in prosthetic valve dynamics was initially stimulated during formation of the Cardiac Research Development Laboratory tasked with supporting the newly opened Cardiac Surgery unit at Royal Jubilee Hospital in Victoria, BC, Canada (1973). Incorporated as Vivitro Systems Inc (VSI), our primary focus was research, design, and development of cardiac valve implant devices and on the laboratory test systems required. During this phase we studied valve motion in an early pulse duplicator using high-speed cinematography and photogrammetric analysis (Brownlee and Scotten 1976; Scotten et al. 1979). Subsequently, an innovative simpler method was devised to determine projected open area (POA) like planimetric quantification of POA from 16 mm cinematography and was published in *New Laboratory Technique Measures Projected Dynamic Area of Prosthetic Heart Valves* (Scotten and Walker 2004). In 2009, work transitioned into a separate independent research and development enterprise, ViVitro Laboratories Inc. (VLI) also based in Victoria, BC, Canada.

Consistent with a focus on evaluation of various prosthetic valve models, our pulse duplicator was modified to include a unique opto-electronic subsystem which we named *Leonardo*. In addition to developing this subsystem, we shifted focus to heart valve closure phase dynamics and conspicuous supra-physiologic backflow fluid velocities. Driven by ongoing curiosity and armed with new data from *Leonardo*, we reported our findings through a series of preprints and publications (Scotten and Walker 2004; Scotten et al. 2006, 2011, 2014, 2015 preprint versions 1-4, 2019-2020, 2022; Chaux et al. 2016; Deutsch et al. 2018; Lee et al. 2020; Lee et al. 2021). In a recent broader application of our investigative technology, results from *in silico* computational modeling and *in vitro* experimental studies confirmed or verified the characteristics of leaflet spatial oscillations in bioprosthetic valves (flutter) throughout the open period of the cardiac cycle Lee et al. 2020 and prompted peer commentary (Lee et al. 2021; Carpenter 2021; Obrist and Carrel 2021; Carrel et al. 2023; Barret et al. 2023).

Given the arc of success with prosthetic valves over decades including initiation and progressive expansion of transcatheter devices, long-term durability and thrombosis issues persist. Thrombosis is a physics-based phenomenon that nature evolved to stem bleeding after an injury. For both transcatheter and surgically implanted bioprosthetic valves, limited durability related to multiple factors has stimulated introduction of a variety of “rescue devices”. Intended to provide transcatheter based mitigation of complications in primary and valve-in-valve bioprosthetic valves implants, these devices are associated with their own unique complications. The longer-term consequences of a transcatheter based multiple valve approach for patient morbidity, mortality and overall cost are yet to be determined. In the current paper, our focus returns to identification and assessment of sources of thrombogenicity in contemporary clinical and experimental mechanical and bioprosthetic heart valves with particular attention to the central role of transient fluid velocities during valve closure. Flow velocity manifests shear via flow velocity gradients that can trigger blood damage and clot formation in vascular disease processes and cardiovascular implant devices (Obrist and Carrel 2021; Chan et al. 2022; Sheriff et al. 2021). When adjacent fluid layers bypass with speed differentials, shear forces increase and blood damage results. We have sought to assess and compare the dynamic behavior of clinical and experimental valves which stimulated provocative conclusions regarding development of less thrombogenic devices.

## METHODS

### Pulse Duplicator Experiments and Computational Modeling

Over time, progressive adaptations to the pulse duplicator and experiment outcomes have been reported (Lee et al. 2020; Lee et al. 2021; Scotten et al. 1979; Scotten and Walker 2004; Scotten et al. 2006; Scotten and Siegel 2011; Scotten and Siegel 2014; Scotten and Siegel 2015; Scotten et al. versions 1-9, 2019-2020). This included the optical measurement of valve POA kinematics and non-trivial *in silico* evaluations (Barret et al. 2023; Kovarovic et al. 2023; Lee et al. 2020; Lee et al. 2021; Scotten et al. 2022). Test conditions, FSI parameters, and boundary conditions used in this study are reported in the Fig. 1 caption. Computational model results and experimental outcomes are in close agreement (Lee et al. 2020; Lee et al. 2021).

**Figure 1.**
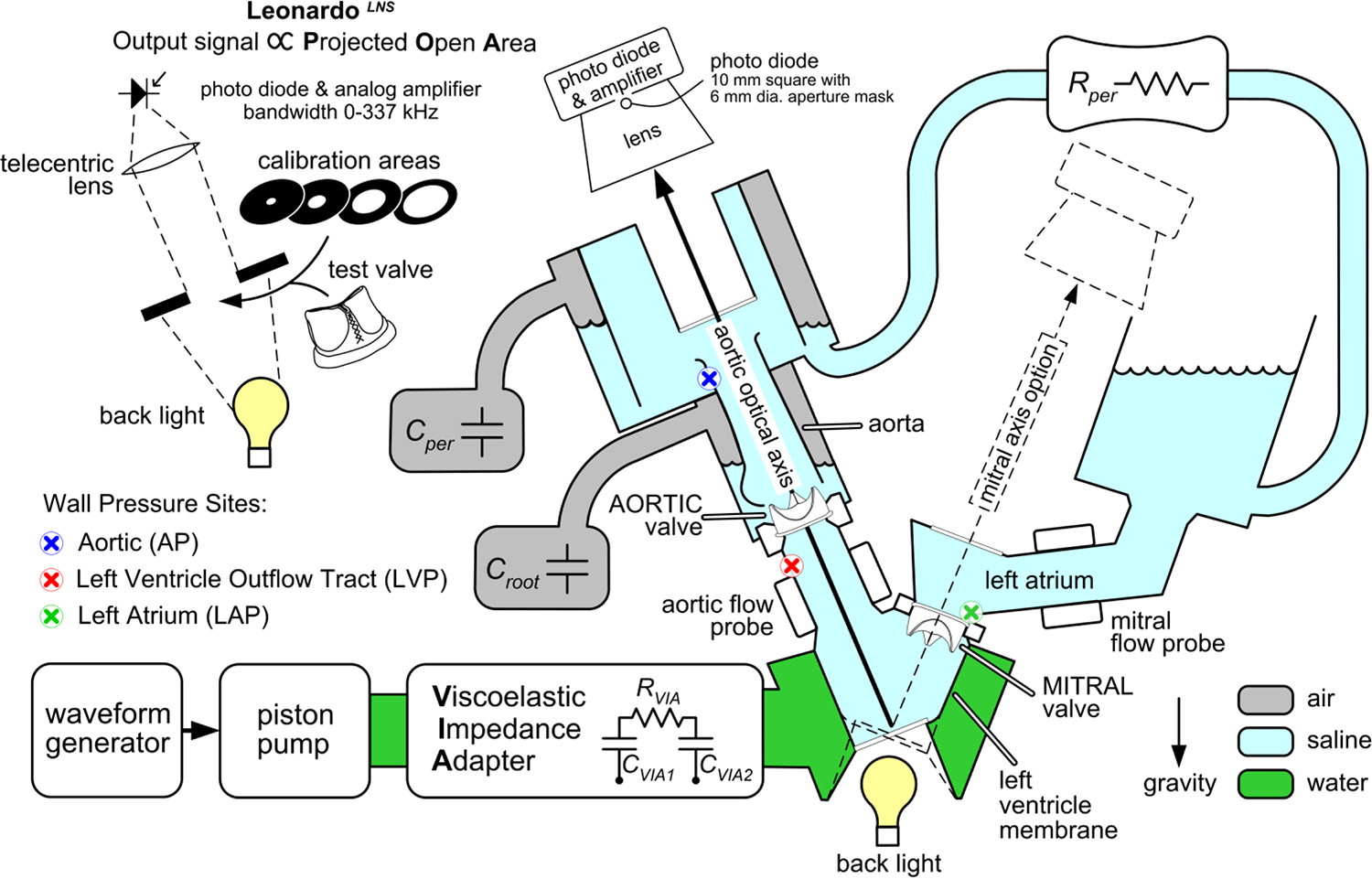
Leonardo*^LNS^* pulse duplicator system has reduced models of *C*ompliance and *R*esistance: *C_VIA1_* 120 mL; *C_VIA2_* 50 mL; *C*_per_ 615 mL; *C_root_* 640 mL *R*_per_ adjusted for normal mean aortic pressure of 100 mm Hg; *R*_VIA_ 200 c.g.s. units test fluid saline; cardiac output ∼ 5 L/min

**Figure 2.**
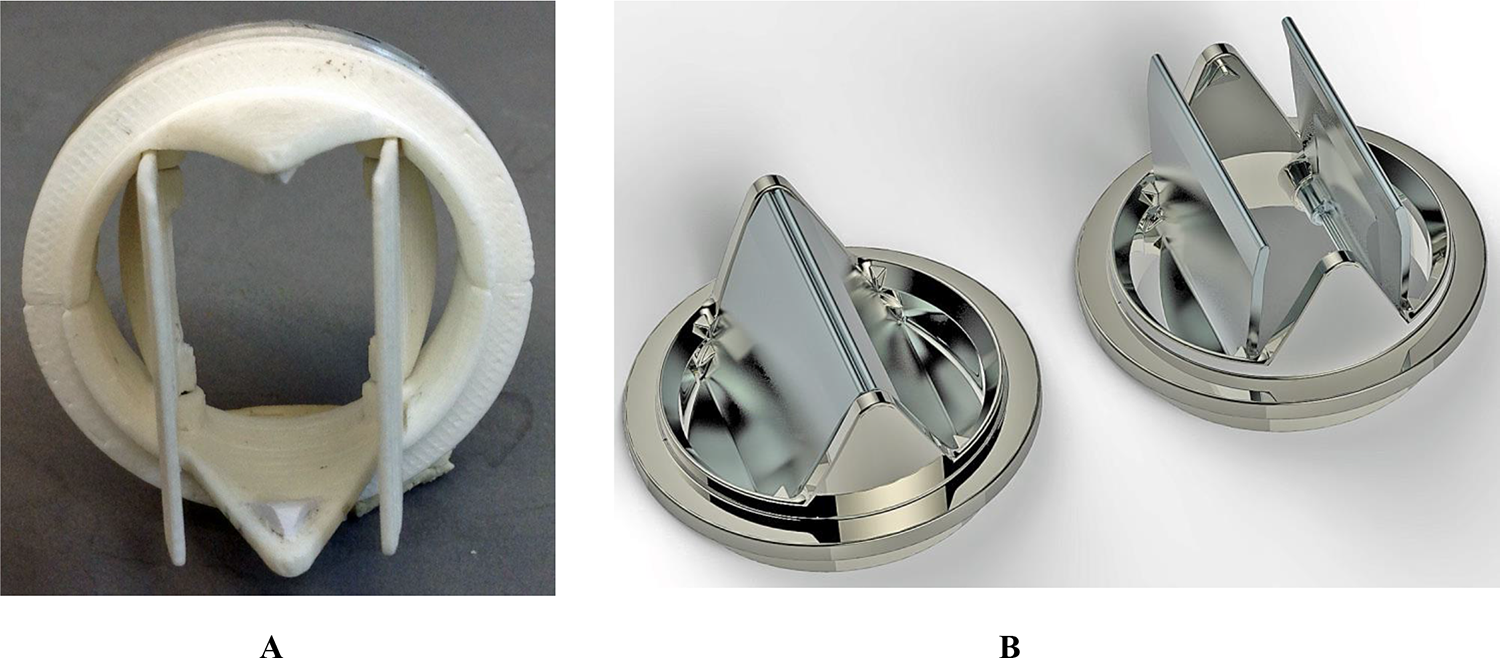
(A) photograph (B) rendering of prototype bi-leaflet mechanical valve BV3D See video https://youtu.be/5QmDPbfUvTManimation_link

Test valves and controls are listed in Table 1. Flow and pressure signals are filtered by analogue circuitry (bandwidth BW∼ 0-100 Hz). The POA signal was unfiltered had a rise time ∼ 1.04 µs (BW ∼ 0-337 KHz). Saline used is an accepted test fluid per ISO 5840-3 with density ρ=1.0 g/cm^3^ and dynamic viscosity µ=1.0 cP (Cardiovascular Implants –cardiac valve prostheses –part 3: ISO 2013).

**Table 1.** Test valves and controls.

| Valve Name | Description | (A)ortic, (M)itral | Seat Size (mm) |
| --- | --- | --- | --- |
| <sup>1</sup> St. Jude Medical (SJM) Regent<sup>TM</sup> | Mechanical bileaflet | A, M | 25 |
| mock-TAVR | Simulated TAVR with preset trivial para-valvular leak | A | 25 |
| <sup>2</sup> On-X<sup>TM</sup> | Mechanical bileaflet | A, M | 25 |
| BV3D<sup>LNS</sup> | Printed prototype, bileaflet | A, M | 25 |
| <sup>3</sup> Lapeyre-Triflo<sup>TM</sup> -F6 (circa 2004) | Prototype, mechanical tri-leaflet (carbon occluders) | A, M | 25 |
| <sup>4</sup> Novostia Triflo PEEK (circa 2023) | Mechanical tri-leaflet (PEEK occluders) | A | 21 |
| <sup>5</sup> Edwards-Perimount<sup>TM</sup> | Bioprosthetic, pericardial tri-leaflet | A, M | 25 |
| <sup>6</sup> Sorin Mitroflow<sup>TM</sup> | Control pericardial tri-leaflet bioprosthetic | A | 21 |
|  | (Non-test) pericardial tri-leaflet bioprosthetic | A/M | 29 |

<sup>1</sup> Abbott Laboratories, Chicago IL, USA. <sup>2</sup> Artvion Inc., Kennesaw, GA, USA. <sup>3</sup> Lapeyre-Triflo<sup>TM</sup> -F6, <sup>4</sup> Novostia-Triflo PEEK, <sup>5</sup> Edwards-Perimount<sup>TM</sup>, Edwards Lifesciences, Irvine, CA, USA, <sup>6</sup> Mitroflow Sorin, Salluggia, Italy.

### Assessment of dynamic flow velocity

The Leonardo system enables assessment of dynamic flow velocity through test valves using measurements from an electromagnetic magnetic flow probe together with photodiode detection of POA. Mathematically, flow velocity equates to the ratio of these two quantities as:

*flow velocity (cm/s) = volumetric flow rate (cm^3^/s) / POA (cm^2^)*

The assumption is POA has a uniform flow profile, but this may not hold true when valve occluder motion and flow are irregular. Experiments indicate that as POA decreases, valve flow velocity increases and conversely, decreases when POA increases. This behavior is governed by Bernoulli’s principle.

### Pressure measurement

Left atrial, left ventricle aortic outflow tract, and aortic sites pressures measured via short catheters (∼ 7.5 cm length) have internal diameter of ∼1.8 mm connected to disposable pressure transducers (Deltran^TM^, model PT43-604)**. Catheter ports measured wall pressures referenced to the mid-plane of the test valve. In Fig. 1, aortic transvalve pressure is measured between the left ventricular outflow tract (LVP) and the aorta (AP).

Mitral transvalve pressure is measured between the left atrium (LAP) and the left ventricular outflow tract (LVP). A TriPack TP2001 unit *(ViVitro Systems Inc., circa 2001*) contains three differential amplifiers with switch selectable low-pass filters (AM9991 plug-in boards).

### Significant waveform characteristics

Several traits are clear in Figs. 3-5. All signals were sampled synchronously. Regions relevant to the valve closure moment are near the dashed red line. Of importance are initial minimum valve POA values and initial peak negative values attained in transvalve pressure, volume flow rate, and closure flow velocities. For bioprosthetic valves, we observed an upward-downward movement of both the valve frame and leaflets demonstrative of compliance reactivity. Hydrodynamic oscillations are also present post valve closure as seen in the unadjusted volume flow rate in some of the waveforms. Comparing the phasing of volume flow rate and POA near valve closure, regional components having compliance influence hydrodynamic patterns.

**Figure 3.**
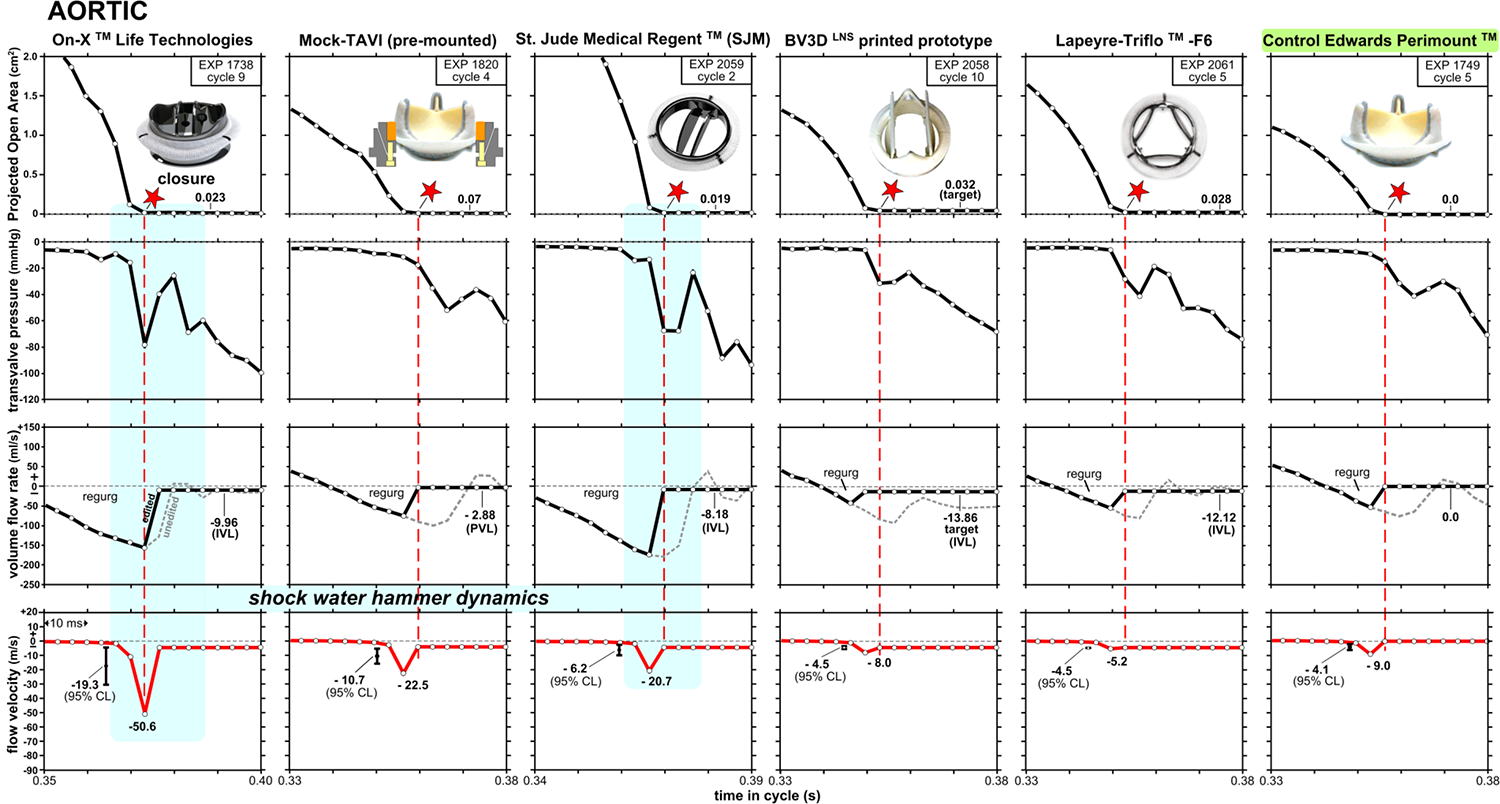
AORTIC valve test data includes 10 consecutive cycles for each experiment. For specified cycles, valve flow velocity (red waveforms) equates to volume flow rate/ POA. Confidence limits (CL) illustrate range for 95% of flow velocities. Note: peak flow velocities may fall outside the bounds of CL. Data sampling interval of the white dots is 3.36 ms. Intra-valvular leakages are denoted (IVL) and para-valvular (PVL). Edited *vs.* unedited volume flow rate waveforms are displayed, respectively, as solid black and dashed grey. Flow velocity profiles (red) negate the regional reactive dynamics associated with elasticity and inertia. Shock water hammer zones are highlighted in blue. Control valve: bioprosthetic Edwards Perimount [25A].

**Figure 4.**
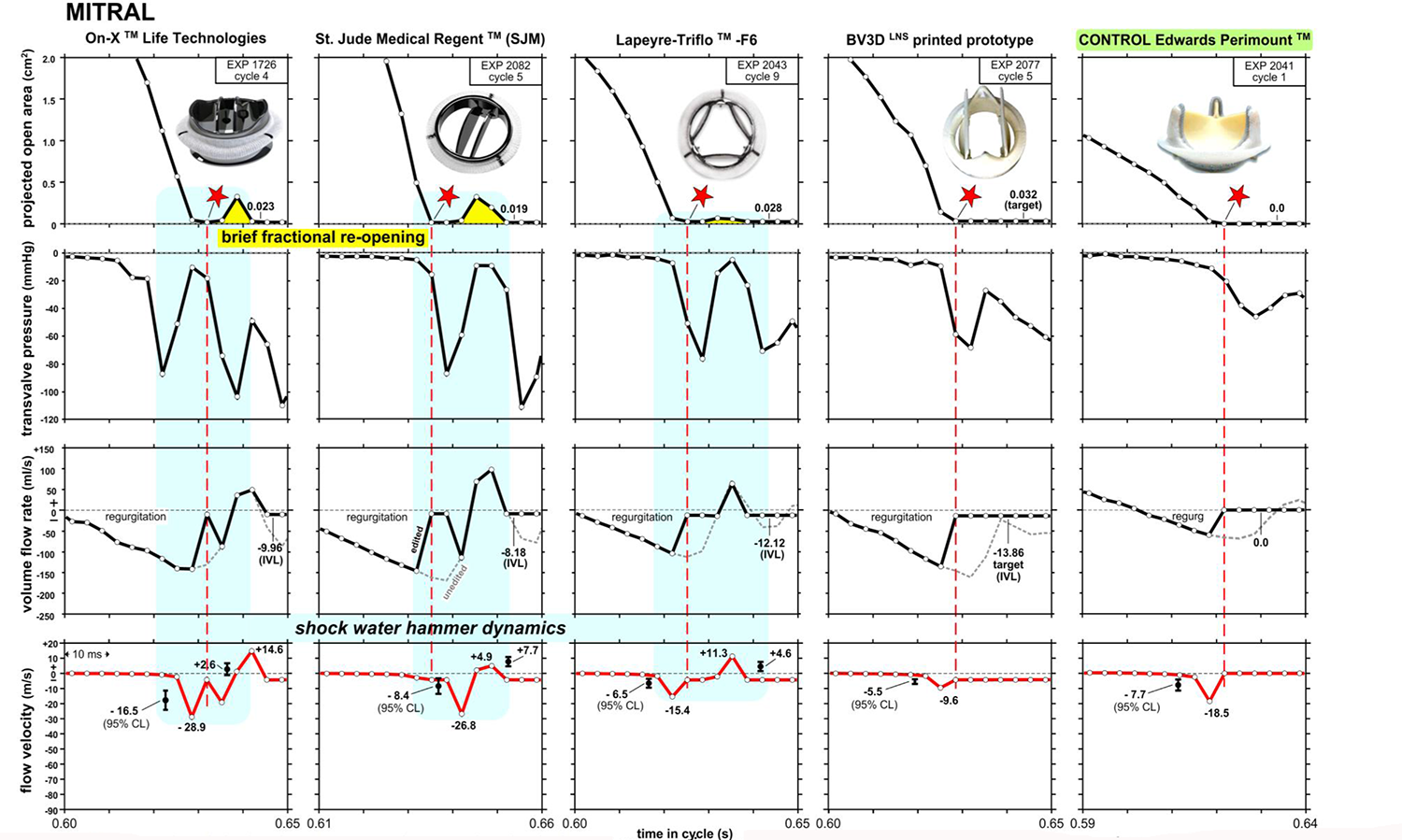
Measured and derived MlTRAL valve test data includes 10 consecutive cycles each experiment. A spatially averaged metric for mitral valve flow velocity (red curves) is equated for specified cycles to volume flow rate/ valve projected open area. Mean flow velocity and 95% confidence limits are shown. Some peak flow velocities shown are outside the upper and lower bounds. An optical approach measures valve projected open area. Data acquisition sampling interval of white dots is 3.36 ms. Intra valvular leakage rates are shown (IVL) and paravalvular (PVL). Edited *vs.* unedited volume flow rates are shown dashed grey alongside edited solid black waveforms. Influence ofreactive elastic and inertia elements are apparent in the unedited flows (dashed grey) which are edited out prior to estimation the red flow velocity waveforms. Partial re-opening of MITRAL projected open areas are shown highlighted in yellow and the shock water hammer zones are in blue. Control valve is Edwards Perimount.

**Figure 5.**
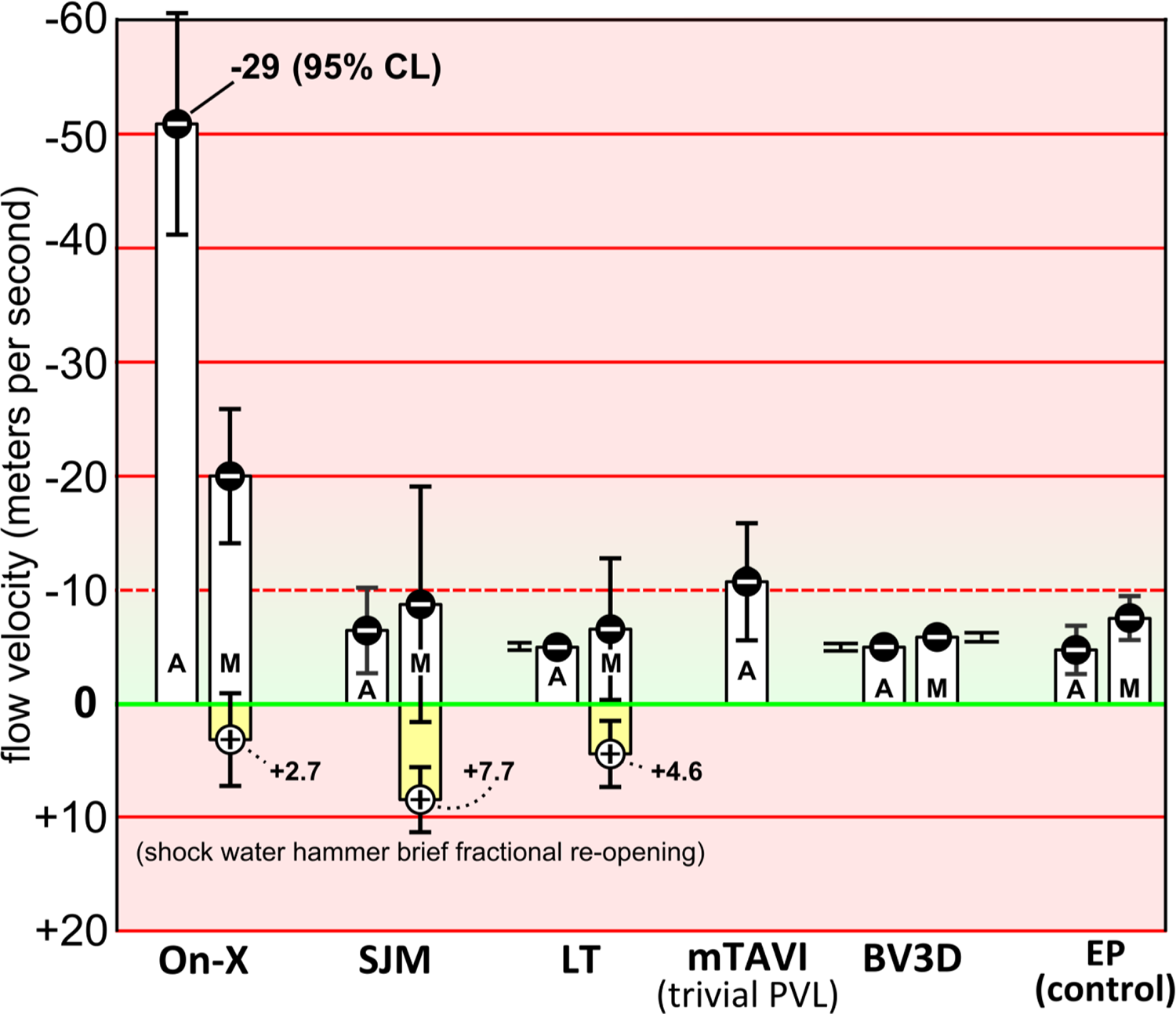
**A**ortic, **M**itral valves are of seat size 25 mm. Average valve closing flow velocity is for ten consecutive cycles. Confidence limits (CL) illustrate the range done for 95% of flow velocities. Valves include: -Mechanical (bi-leaflet: **On-X**, **SJM**, printed prototype **BV3D**), (carbon tri-leaflet design, **LT** - F6), -Tissue (mock-TAVI, **mTAVI**, Edwards-Perimount^TM^ **EP**).

**Figure 6.**
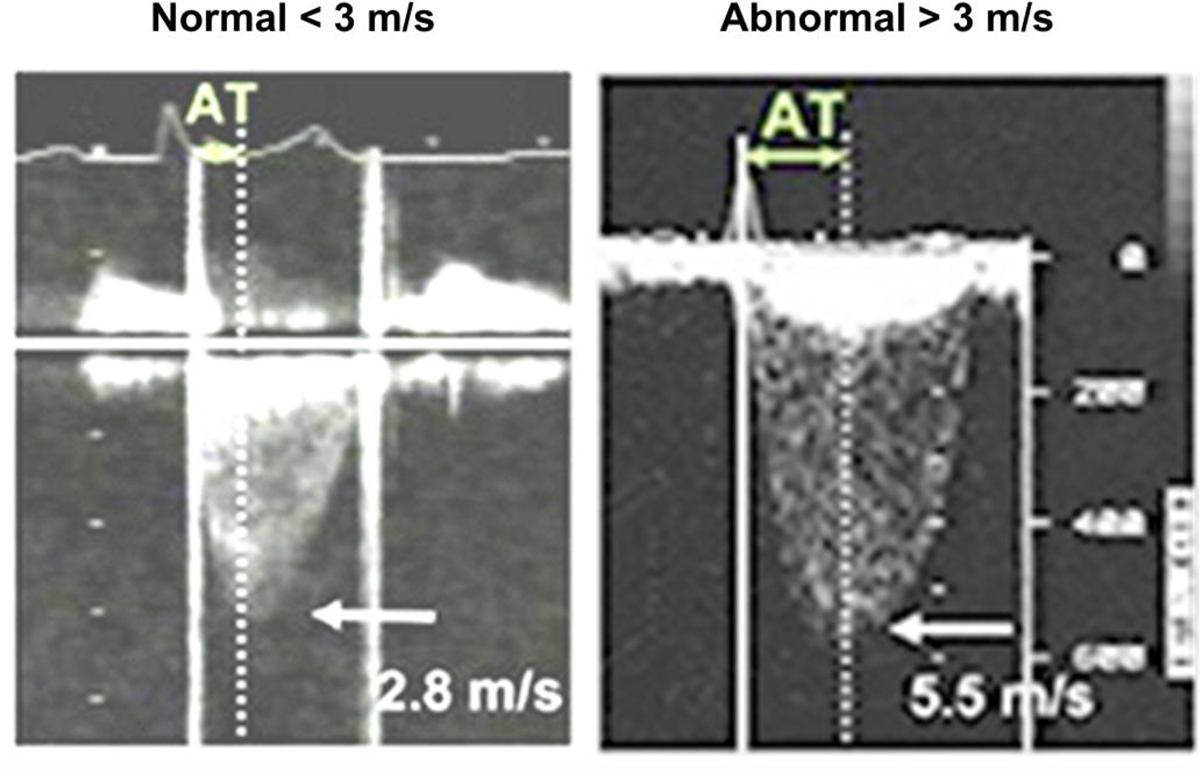
Examples of peak CW Doppler signals showing prosthetic aortic valve forward flow velocity. A peak clinical forward flow velocity of 2.8 m/s is considered normal. <u>Link</u>. Thus, the On-X aortic valve with in vitro flow velocity of 1.8 m/s, affirms normal functioning (see Fig. S2).

**\*\*** Utah Medical Products Inc., Midvale, Utah 84047-1048, USA

**Figure 3.11.**
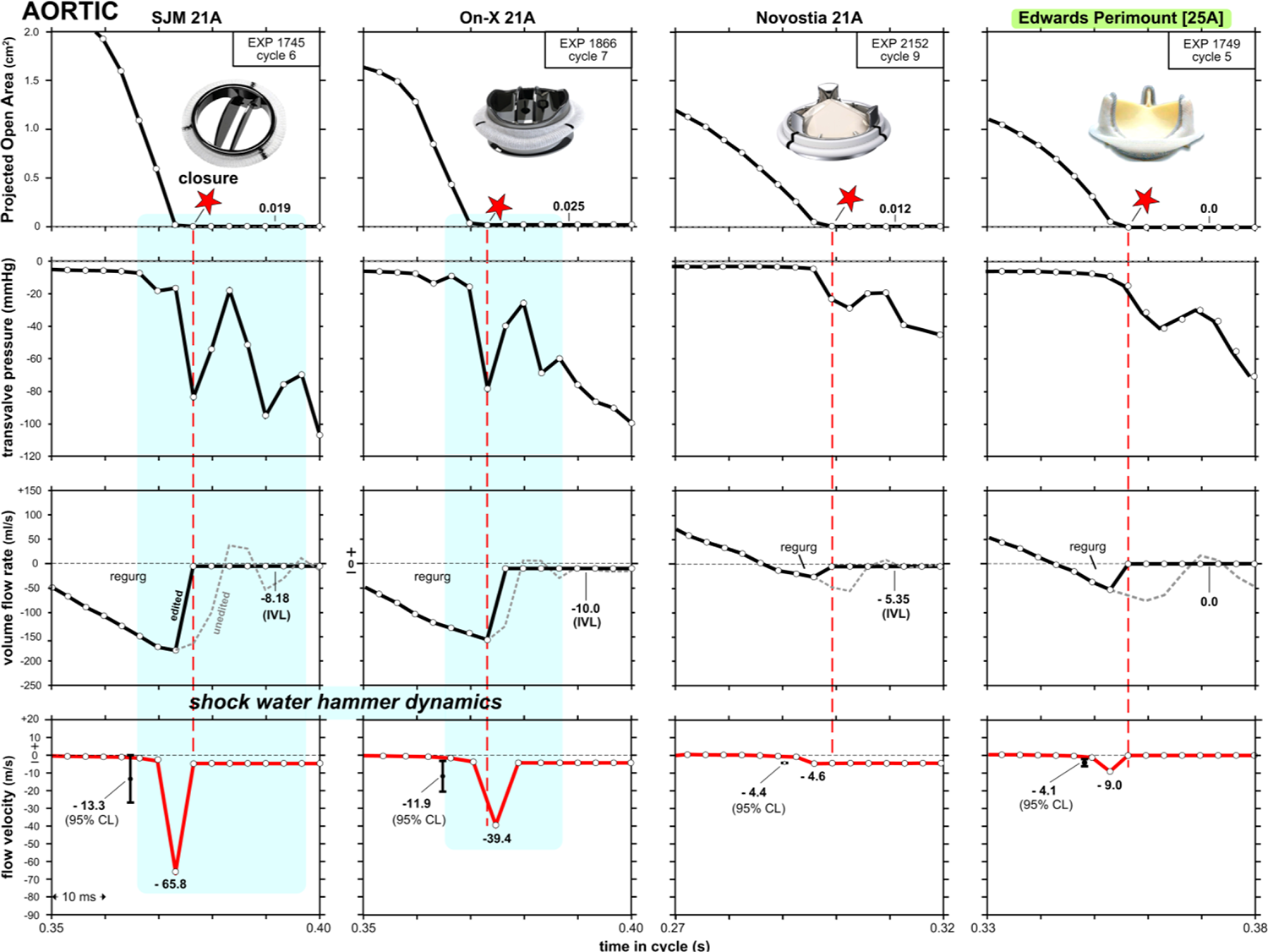
The present-day model of the Novostia mechanical trileaflet valve, size 21 AORTIC (*August 2023*) measures least closing flow velocity compared to the other valves for 10-consecutive cycles. Confidence limits illustrate a range for 95% of flow velocities but peak flow velocities can be outside the bounds of CL. For specific cycles, valve flow velocity profiles (red) equate to volume flow rate **/** POA. Edited *vs.* unedited volume flow rate profiles are solid black alongside the dashed grey profiles. Compliance and inertia reactivity are negated while computing the edited (black) flow rate profiles. Data sampling interval of 3.36 ms is denoted by white data points. Intra-valvular leakages are denoted (IVL). Shock water hammer zones are highlighted in blue. Control valve in this figure is for an Edwards Perimount [25A]. Additional control data for a Mitroflow a size 21A pericardial bioprosthetic valve is provided in supplement Figure S1.1.

### Signal synchronicity

As indicated in Figs. 3 and 4, in the aortic and mitral valve sites, instantaneous minimum regurgitant volume flow rate and minimum POA are synchronous with valve closure. The flow and pressure signals are processed through 100 Hz low-pass analogue filters whereas POA is unfiltered to preserve maximum frequency response (∼ 0-337 KHz).

Synchronized hydrodynamic oscillations emerge post valve closure due to the pooled reactive compliances of the test valve and holder. These damped oscillations are observed in the unadjusted volume flow rate signals (dashed grey) and are ascribed to the combined movement of test valve components (i.e., leaflets plus stent) within the elastic silicone rubber holder. Oscillations evident for all valves and sites have similar periodicity ∼ 20ms (50 Hz).

### Statistical Analysis

For each experiment, we acquired 10 consecutive cycles and reported average velocity measurements and cycle-to-cycle variations of 10 negative peak flow velocities recorded using confidence limits (CL). Summary of the valve flow velocity test datasets utilize an EXCEL*** data analysis tool labeled descriptive statistics together with the options - Analysis Tool Pak and - Solver Add-in. In Figs. 3-5, mean negative peak flow velocities and CL=95% indicated by CL bars adjacent to the flow velocity waveforms show the predicted uncertainty.

*** Microsoft, Redmond, WA, USA

### Experimental valve design

The BV3D rapid prototype valve design shown in Figs. 2A and 2B are a working model that resulted from several designs trialed (Scotten and Siegel 2015). Consideration of competing factors produced 56 laboratory constructed prototype valves with data from experiments producing empiric optimization through interaction of leaflet profile, pivot location and surface characteristics. These led to the rapid printed prototype model BV3D with advantageous forward and especially closing flow dynamics. In this work, *Leonardo* provided immediate feedback on the impact of even subtle geometric changes on valve dynamic behavior.

## RESULTS

### Aortic *vs.* Mitral Dynamics

When fluid in motion is abruptly halted or changes its direction (momentum change) water hammer is observed. This unwanted transient motion closely associates with valve closure timing, property of valve mountings and has been previously reported in vitro (Scotten and Siegel 2015). Figs. 3-5 illustrate snapshots of shock water-hammer dynamics with mean flow velocities reaching −29.0 m/s for the On-X mitral and −65.2 m/s for the On-X aortic valve and is absent for the Lapeyre-Triflo -F6, prototype BV3D, mock-TAVI, and Edwards-Perimount control valves. The mock-TAVI valve was untested in the mitral site.

For the On-X, SJM, and LT valves in the mitral site, a brief fractional reopening is evident in POA post initial valve closure. In Fig. 4, negative peak transvalve pressure spikes range over comparable levels in the mitral site (−75 to −90 mmHg). However, in the aortic position (Fig. 3) pressure spikes encompass a wider range (−40 to −95 mmHg).

Signature shock water hammer dynamics for aortic and mitral valves are evident for POA, transvalve pressures, volume flow rates, and derived valve flow velocity waveforms. We found that shock water hammer dynamics can be mitigated by valve designs optimized for soft valve closure which reduces retrograde valve flow velocity in the early closing phase as observed for the BV3D and EP valves.

### Edited *vs.* unedited instantaneous volumetric flow rates

In Figs. 3 and 4, measured and derived aortic and mitral valve test results include 10 consecutive cycles each experiment. Data acquisition sampling interval is ∼3.36 ms (white data points). Evident are maximum negative flow velocities at valve closure ranging up to −65.2 m/s (On-X aortic). The oscillatory patterns seen in the primary unadjusted (grey) flow rate waveforms were edited out to produce the black flow rate profiles prior to deriving the red flow velocity profiles. The spatial average of mitral flow velocities (red profiles) is obtained by dividing time-periodic volume flow rate by time-periodic POA. Average flow velocity and CL bars show where 95% of the derived negative peak valve flow velocities fall. An inherent advantage of *Leonardo* is that POA is determined by a high resolution opto-electronic methodology with excellent spatial area resolution (details in supplementary information **S2 –** Leonardo sub-system opto-electronics). Leonardo offers clear detection of initial and water hammer rebound events. Distinctive valve motion events such as: initial POA opening, POA closure, shock water hammer rebound where POA goes briefly positive just after valve initial closure are all discernible in the POA waveforms. Volumetric flow rate, transvalve pressures, and derived flow velocity waveforms are acquired synchronously. In Fig. 4, the mock-TAVI valve produced aortic but not mitral site data. Leakage rates are indicated either intra-valvular (IVL) or para valvular (PVL). Highlighted in yellow are post valve closure fractional re-openings attributable to shock water hammer phenomena.

*Readers will note that in prior publications, closing flow velocities more than those observed in this study were reported (Scotten et al. 2004; Scotten and Siegel 2014; Chaux et al. 2016; Deutsch et al. 2018)*. *Previous records with excessive closing flow velocities have been rectified in this report by re-analysis of the historical experiment test datasets imported into revised EXCEL templates*.

The relationship between POA, flow velocity, and mass flow rate helps in understanding the fluid dynamics of valves. Factors such as fluid properties, valve design, and pressure differentials influence this relationship. For example, prototype valves with non-curved leaflet profiles and pivot locations near the orifice center show promise in optimizing valve designs for reduced thrombogenic potential. Partially closed valves evidenced by decreased POA as well as an increase in flow velocity, aligns with the Continuity Principle. The continuity equation states that the mass flow rate of a fluid must remain constant along a continuous flow path and is expressed mathematically as Q = A_1_V_1_ = A_2_V_2_ where Q is the fluid flow rate, A_1_ and A_2_ are the areas of the pipe or valve at two different locations, and V_1_ and V_2_ correspond to velocities of the fluid at those locations. Therefore, when the area available for flow is least (i.e., when the valve is approaching closure and almost closed), the velocity of the fluid is greatest. Transvalve pressure is amplified by increased flow velocity and, in accordance with Bernoulli’s principle, assists valve closure indicated by a reducing POA.

Although high blood flow velocity alone may not cause blood damage directly, the presence of high flow velocity gradients which occur when flow velocity varies significantly over a short distance, can potentially trigger clotting. Such velocity gradients (shear) often arise in areas along the flow pathway where flow disturbances such as constrictions, or turbulence occur. Shear refers to the frictional force between adjacent fluid layers moving at different velocities. Steep velocity gradients contribute to increased shear potential. When shear force exceeds the physiological tolerance, cell fragmentation and activation of clotting processes can follow. While blood cells are capable of withstanding moderate shear levels due to their deformable membranes, certain implant devices like mechanical heart valves are deemed thrombogenic because shear forces expose cells beyond their physiologic limits. Consequently, individuals with such devices may require chronic anti-coagulation therapy to mitigate clotting risks.

Transvalve flow rate is determined primarily by valve open area and transvalve pressure. When valve open area is decreasing (partially closed), transvalve pressure and flow velocity increase. This relationship is governed by Bernoulli’s principle which states that as fluid velocity increases fluid pressure decreases. Therefore, when POA is decreasing (valve not completely closed), fluid velocity increases, transvalve pressure increases and open area declines. Conversely, when the valve is fully open, and pooled open areas are maximal, fluid velocity declines in conjunction with transvalve pressure resulting in lower volumetric flow rate. High blood flow velocity may not be an independent trigger for blood damage, but generation of <u>high flow velocity gradients</u> can potentially lead to clotting activation. Velocity gradients come about when there is a significant change in flow velocity over a short distance and are often observed in areas having flow pathway disturbance, constriction, or turbulence.

Shear is a frictional force exerted by adjacent fluid layers moving at different velocities. With steep velocity gradients, shear potential increases. When physiological tolerance for shear is exceeded, cell fragmentation and activation of clotting processes may result. While blood cells have deformable membranes that can tolerate moderate levels of shear, certain implant devices, such as mechanical heart valves, can be overtly thrombogenic when exposed to high shear force requiring chronic anti-coagulation therapy for recipients.

Several notable points in Figs 3-5 are:

1. The occurrence of positive flow velocities shortly after valve closure for the SJM, On-X and LT.
2. The On-X, SJM, LT mechanical mitral valves show shock water-hammer patterns having momentary post closure fractional reopening.
3. Shock water-hammer patterns are absent for the two bioprostheses (EP and mTAVI).
4. The EP aortic and mitral control valves exhibit low closure flow velocity (< 10 m/s).
5. For the SJM valve, the differential between aortic and mitral maximum volumetric backflow rates consistently exceeded that observed in other tested valves ∼ (−180 ml/s _-max_ *vs.* −150 ml/s _-max_).
6. For current clinical mechanical mitral valves, a disquieting abrupt closure and shock water hammer dynamics is observed and shown highlighted in yellow. In this region, a brief partial valve re-opening is triggered ∼ 6 ms after valve closure with rebound duration of ∼ 4-10 ms.
7. Mitral valve closure generates oscillations in flow rate and transvalve pressure greater in amplitude but less dampened than valves in the aortic site.
8. Mitral valve occluder rebound (highlighted yellow) associates with high negative regurgitant volume flow rate, transvalve pressure peaks and flow velocities which impacts on valve closing with certainty.
9. Negative transvalve pressure spikes are near synchronous with valve closure.
10. Transvalve pressure phase leads the flow.
11. Important features underscored in the valve closure phase are near synchronous timing of valve closure (i.e., valve minimum POA), negative going transvalve pressures at closure, maximum retrograde closure volume flow rate, and maximum retrograde flow velocities.
12. The velocity of unsteady pressures in water is ∼1,500 m/s. Therefore, pressures in immediate proximity to the aortic test valve would, for example, have trivial phase (∼0.07 ms) relative to the aortic wall pressure site which is ∼10 cm distant from the mid-plane of the valve.

Results in Figs. 3-5 are based on predicted valve flow velocities. Notable near post valve closure are <u>positive</u> <u>flow velocities</u> for SJM, On-X and LT. An interesting mitral site pattern shows shock water-hammer phenomena associated with three of the four mechanical mitral valves tested (On-X, SJM, LT), where kinematic evidence shows momentary post closure fractional reopening and absence in the two bioprostheses (mTAVI and EP). The EP aortic and mitral control valves have low closure flow velocity (< 10 m/s).

Important elements in Figs. 3-5 are:

1. The EP aortic and mitral control valve demonstrated the lowest predicted valve closing flow velocity which is consistent with their reported clinical experience of low thrombogenic potential.
2. Driven by shock water hammer dynamics, backflow velocity is higher in the mitral than aortic site.
3. Slow-motion and real time visualization of aortic and mitral valves qualitatively revealed both symmetrical and non-symmetrical closing behavior of occluders. Quantitatively, flow velocity confidence levels (CL) may be a useful measure of occluder motion evenness over 10 consecutives cycles. Results show that the On-X and SJM have greatest CL values.
4. The mTAVI valve sample with trivial para-valvular leak (PVL) had a peak average negative valve closing flow velocity of −10.7 m/s (±5.25). However, it has been noted that valve samples with PVL = zero and counter intuitively, with > PVLs, have reduced valve closing flow velocities (Scotten and Siegel 2011; Scotten and Siegel 2014).
5. Based on the control valve results, mean closure flow velocities < −10 m/s suggests potential for low/non-thrombogenic shear force.
6. Valve closing dynamics are associated with shock water hammer are also associated with maximums in flow velocity, backflow rate and transvalve pressure.
7. During valve closure, platelets subjected to jet flow velocity gradients are likely to accumulate more shear force damage relative to that from the forward flow phase (Herbertson et al. 2011).

## DISCUSSION

Over the past 60 years, heart valve design and performance evolved to provide improved durability and hemodynamic function. While preferences for mechanical *vs.* bioprosthetic valves fluctuated widely, the advent of transcatheter delivered devices and their less invasive insertion methodology, the balance shifted progressively in favor of bioprostheses. Although initially restricted to use in the elderly or patients designated too fragile for conventional surgical implantation, use is now extended to younger and lower risk candidates.

Younger patient cohorts receiving bioprosthetic heart valves have longer life expectancies (>20 years) and thus a higher likelihood of re-intervention. The shift in favor of BHVs may prove to be premature as more (or lack of) long-term data is in hand (Briffa and Chambers 2017). A recent surgical AVR study by (Lu et al. 2023) in patients aged 50 to 69 years, found long-term survival was better in those who received mechanical compared to Perimount bioprosthetic valves and suggested a substantial survival advantage be recognized in patients with mechanical valves. To respond to an incidence of technical failure in contemporary transcatheter valves and perhaps in anticipation of increasing frequency of degenerated valves in younger patients, a variety of transcatheter “rescue” devices are offered. Inserted within a degenerated or malfunctioning primary valve or a previously implanted rescue device, the predicted durability of rescue devices is speculative. In a prophetic article *Russian Dolls, when valve-in-valve implantation is not enough* (Tseng 2016), Tseng called attention to the possibility of rescue valve recipients who will represent persistent therapeutic challenges that may not be resolvable by transcatheter devices. As a response to a possible increase in this initially small subset of patients, for those valve replacement candidates who favor the alternative of one valve durable for life but are hesitant because of concerns over anticoagulant related issues and the as-yet unfulfilled need by rheumatic patients in developing countries (Zilla et al. 2008; Wium 2020; Wium et al. 2020) it is essential that research driven by purposeful curiosity continues toward achievement of a long awaited anticoagulant independent mechanical valve by research endeavors such as recently reported by (Wium 2020; Wium et al. 2020)

At or near valve closure, flow dynamics can be considered analogous to transitory valve stenosis, whereby regurgitation is increasingly constrained until complete, motionless, closed-valve conditions arise. Thus, during brief crucial moments preceding and after valve closure, localized prothrombotic microenvironments may be relevant to generation of jet flow velocities with shear sufficient to induce blood element damage (Soares et al. 2013; Ding et al. 2015). These influences may impact multiple valve types with potential consequences that may include but not limited to:

- reduced valve leaflet mobility, sub-clinical thrombosis,
- potential for pannus formation,
- cavitation and clinically detectable high intensity transcranial signals (HITS),
- transient ischemic attack (TIA), embolic acute/sub-acute stroke, other silent micro-infarction, and adverse cerebrovascular events (Woldendorp et al. 2021)

### Shock water hammer phenomena and occluder rebound

Mitral valve closure dynamics shown in Fig. 4 include a partial post closure transient opening (rebound) attributable to shock water-hammer, were found less prominent in the aortic site (Fig. 3). Occluder rebound driven by water-hammer power (product of transvalve pressure and volumetric flow rate) for the SJM mitral valve is observed as a momentary post closure partial re-opening. This has been previously reported with high resolution magnified examples of POA rebound data (Scotten and Walker 2004; Scotten and Siegel 2011; Scotten and Siegel 2015). Relative to the bioprosthetic control valve, mechanical mitral valves (On-X, SJM, LT) are driven by higher-level transvalve pressures and retrograde flows and reveal rebound behavior prior to final closure. Additional pro-thrombotic aspects may be related to sub-optimal valve forward flow energy losses (e.g. arterial sclerosis) and biomechanical and biochemical responses sufficient to exacerbate risk of pathologic thrombus formation and propagation.

### Research outcomes reported

(Scotten et al. 2006; Scotten and Siegel 2011; Scotten and Siegel 2014; Scotten and Siegel 2015; Scotten et al. 2020; Lee et al. 2020; Lee et al. 2021; Scotten et al. 2022)

1. High amplitude valve closing flow velocities may result in supra-physiologic shear forces originating in small intra- and para-valvular leak gaps and which become mechanistic initiators of the clotting cascade.
2. Platelet activation is induced by high shear even with short exposure time (Ding et al. 2015).
3. Closing dynamics disparity between mechanical and bioprosthetic valves is chronically overlooked as a primary indicator of valve thrombogenicity.
4. As with transient sparks that can initiate a fire, MHVs exhibit overt closing spikes in flow velocity related to occluder non-response to flow deceleration and residual leakage areas when closed.
5. For a given volumetric backflow rate near valve closure, the smaller the total residual leakage area, the greater the valve closing flow velocity.
6. The highest valve closing flow velocity was for the On-X and the SJM valves compared to tissue control valves (Edwards pericardial 25 A and M, and Mitroflow 21A).
7. The prototype BV3D valve had the lowest predicted valve closing flow velocities in both aortic and mitral sites. Assessment of this dynamic behavior represents a practical means to qualitatively screen valves and controls for thrombogenic potential.
8. Valve BV3D test data suggest that specific MHV leaflet geometries generate a closing force during forward flow deceleration and prior to flow reversal, a potentially beneficial “soft closure” response.
9. Our novel laboratory methodology permits inference of shear damage to formed blood elements and potential for continuing thrombogenic response. This constitutes a mechanistic explanation for observed thrombogenic disparity in prosthetic valve types and designs, now broadened to include observations of TAVR related thromboembolic events.
10. For MHVs, cyclic valve closing flow velocities appear to be related to a prothrombotic state requiring chronic anti-coagulation.
11. Bioprosthetic valves appear not to generate a pro-thrombotic closing phase.
12. In both experimental and computational pulse duplicator studies, bioprosthetic valves in smaller diameters and/or with thicker leaflets generate higher flutter frequencies (Lee et al. 2020; Lee et al. 2021).

### Strengths

A derived valve closing flow velocity provides a useful laboratory metric directly proportional to blood cell damage from shear force. For some mechanical mitral valves, a brief positive flow transient is noted. Such transients result from shock water-hammer dynamics and are indicative of high shear force, blood cell damage, biomechanical and biochemical responses that promote pathologic thrombus formation and propagation.

### Limitations

Mechanical valves tested in the mitral position consistently manifested leaflet rebound observed as a transitory post closure partial re-opening driven by water-hammer power (product of transvalve pressure and volumetric flow rate). This has been previously reported with magnified examples of high resolution POA rebound data (Scotten and Siegel 2014).

To validate computational simulations, high fidelity amplitude and frequency responses are required to resolve small scale valve geometries and hydrodynamic features. This prerequisite may benefit from recently developed computational approaches and extensions needed to help fill lingering computational gaps (Kolahdouz et al. 2020; Kolahdouz et al. 2021).

## CONCLUSIONS

This work exposes the central relationship between thrombogenicity and predicted high velocity flows and shear forces at valve closure. As practical application of our findings, specific valve geometric features were identified that led to prototype designs with potential for further development. The application of unique technology, rapid prototyping and ranking of flow velocity patterns near valve closure optimized experimental valve geometry for reduced thrombus potential compared with control valves.

We compared the hydrodynamics and kinematics of prosthetic heart valves at valve closure to control valves. This data could be predictive of thrombogenic outcomes which is the potential for clot formation. The in-vitro results suggest that the potential for blood clots caused by high velocity flows and shear forces can be reduced by focusing on specific valve geometric features, leading to the development of improved clinical devices. The study however raises the question of whether optimum mechanical valve performance requires similar, identical, or lower valve closing flow velocities compared to contemporary clinical bioprostheses. The work opens an experimentally assisted pathway for developing new, durable and less thrombogenic devices and the prototype model BV3D serves as evidence that focused laboratory efforts can yield promising results.

Additionally, the study indicates that valve flow velocity differentials and associated shear are significant activators of thrombogenicity for the On-X valve, emphasizing the importance of valve closure behavior in reducing thrombus potential and improving clinical devices, despite the challenges in introducing new prosthetic heart valves to the market. The Holy Grail goal of a mechanical valve independent of chronic anticoagulation still beckons.

## Contributions

I. Methodology, conception and design: LN Scotten; (II) Project administrative support: LN Scotten; (III) Provision of study materials: LN Scotten; (IV) Collection and assembly of data: LN Scotten; (V) Data analysis, interpretation, validation: LN Scotten, DJ Blundon; (VI) All authors have contributed.

## Acknowledgements

David Walker is thanked for restoring the software functionality of the PC XP based data acquisition system. Competing Interests: None declared. Financial Disclosure: This research has been done on a pro bono basis by all authors with no financial support of others.

## Data availability statement

The core data that support the findings of this study are available from the corresponding author, [LNS], upon reasonable request.

## Open access

Anyone can share, reuse, remix, or adapt this material, providing this is not done for commercial purposes and the original authors are credited and cited.

## CONTINUES WITH SUPPLEMENTAL MATERIAL…

**Figure S1.1.**
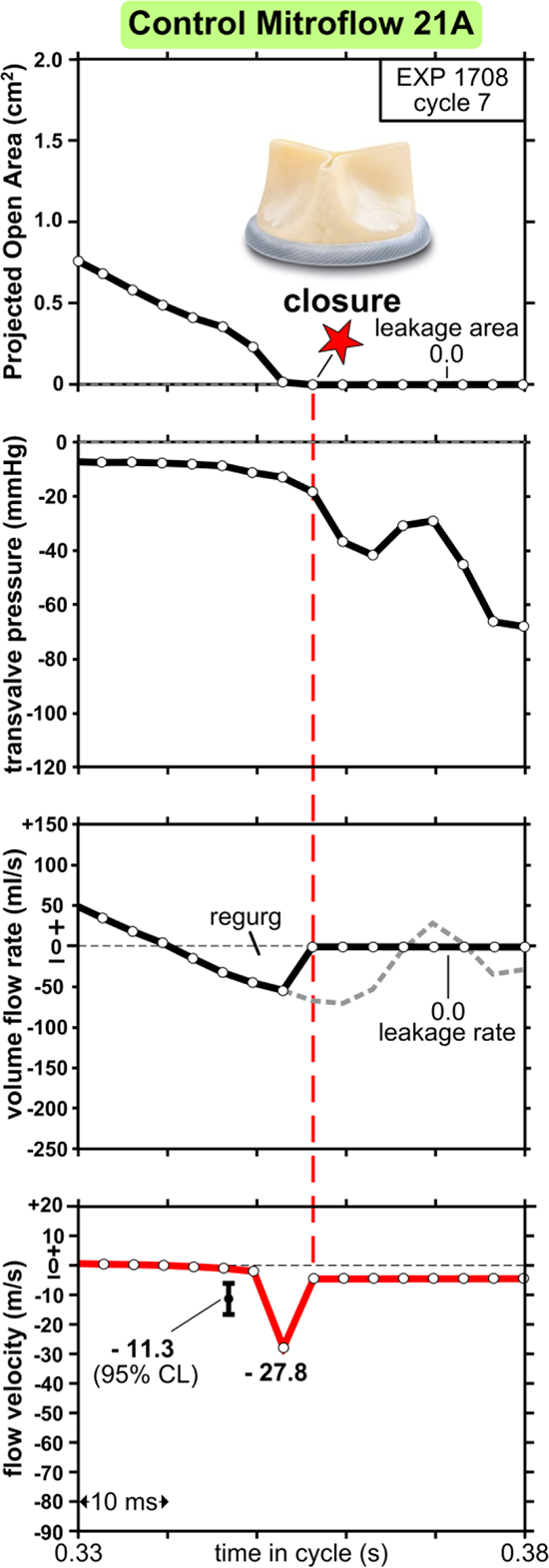
Mitroflow pericardial 21A control valve performance compared to mechanical valves of same seat size. Valve flow velocity (red curve) equates to volume flow rate / projected open area. Mean flow velocity and 95% confidence limits (CL) are shown for 10-consecutive cycles. Note: For cycle 7 a peak minimum flow velocity of (minus) 27.8 m/s, is more extreme than the lower confidence limit. Data acquisition sampling interval (white dots) is 3.36 ms. Leakage area and leakage rate are pointed out. Edited *vs.* unedited volume flow rates are, respectively, solid black waveforms alongside the dashed grey. Regional elastic and inertia reactivity, intrinsic within the unedited flow waveform (dashed grey), is edited out prior to determining the (red) flow velocity waveform and utilizes the solid black edited waveform. Control valve: -Edwards Perimount [25A].

### S2 Leonardo sub-system opto-electronics

The POA signal has high resolution and sensitivity. It utilizes circuitry presented in a book by Hobbs (2008). The circuitry suggested for a photo-diode preamplifier is shown in Figure S2.1. A telecentric lens sets magnification to 0.16 times which images backlit test valves to a diameter ∼6 mm (area ∼0.28 cm^2^). The incident light is limited by a black paper mask aperture of ∼7 mm diameter to enhance the linearity of the photodiode’s response.

**Figure S2.1.**
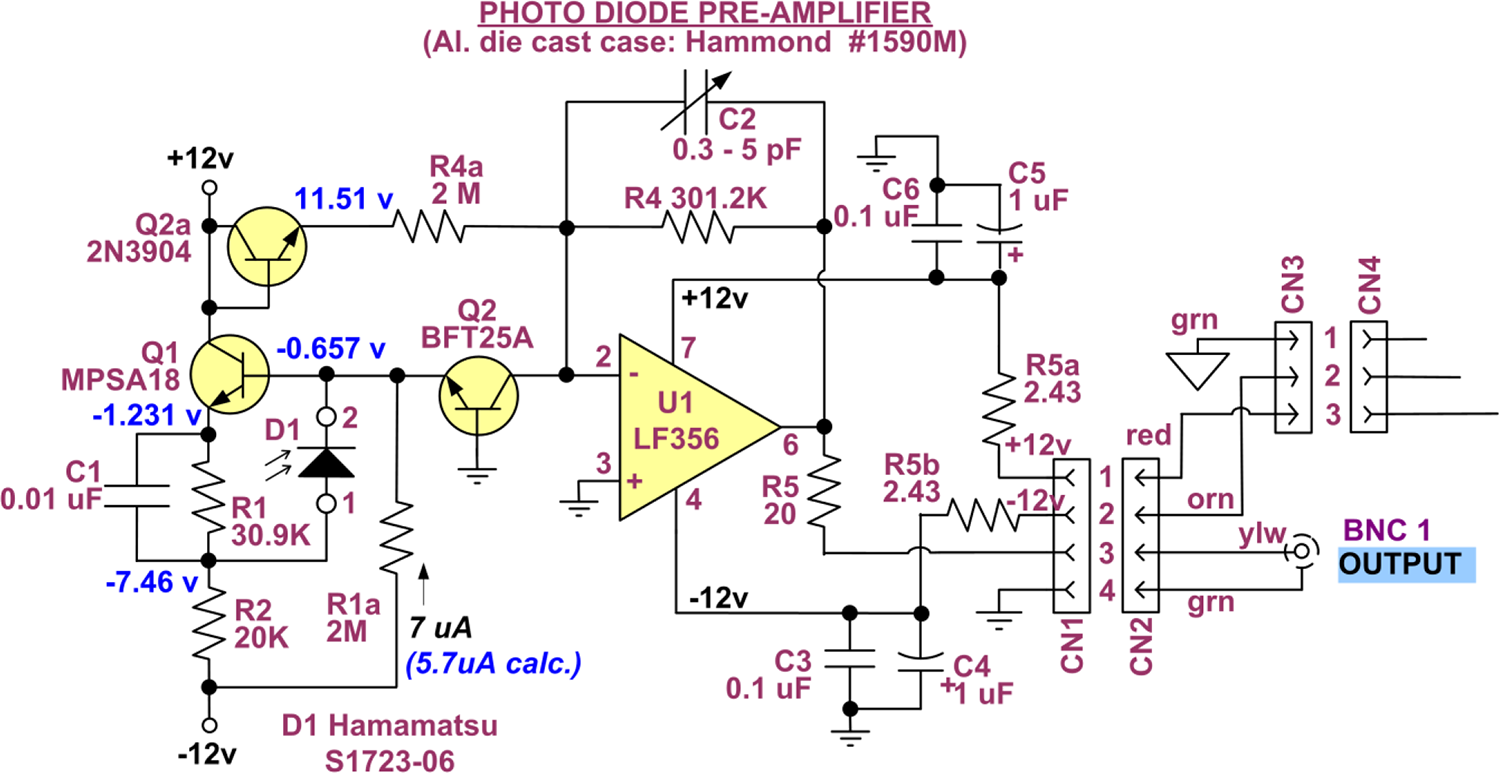
Schematic of photodiode preamplifier shielded by an aluminium die-cast case with tele-centric lens attached via C-mount adapter (below).

**Figure S3.**
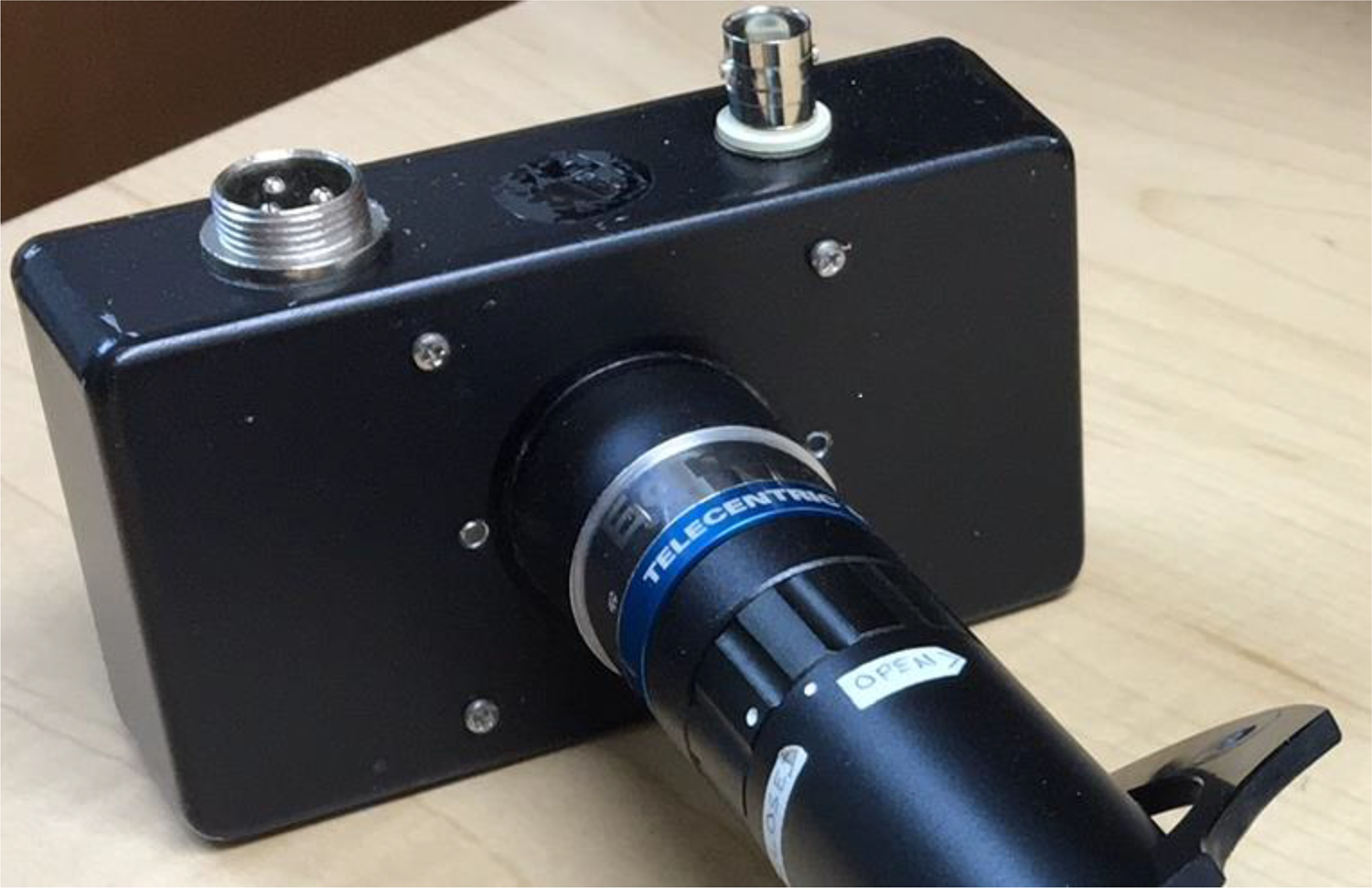
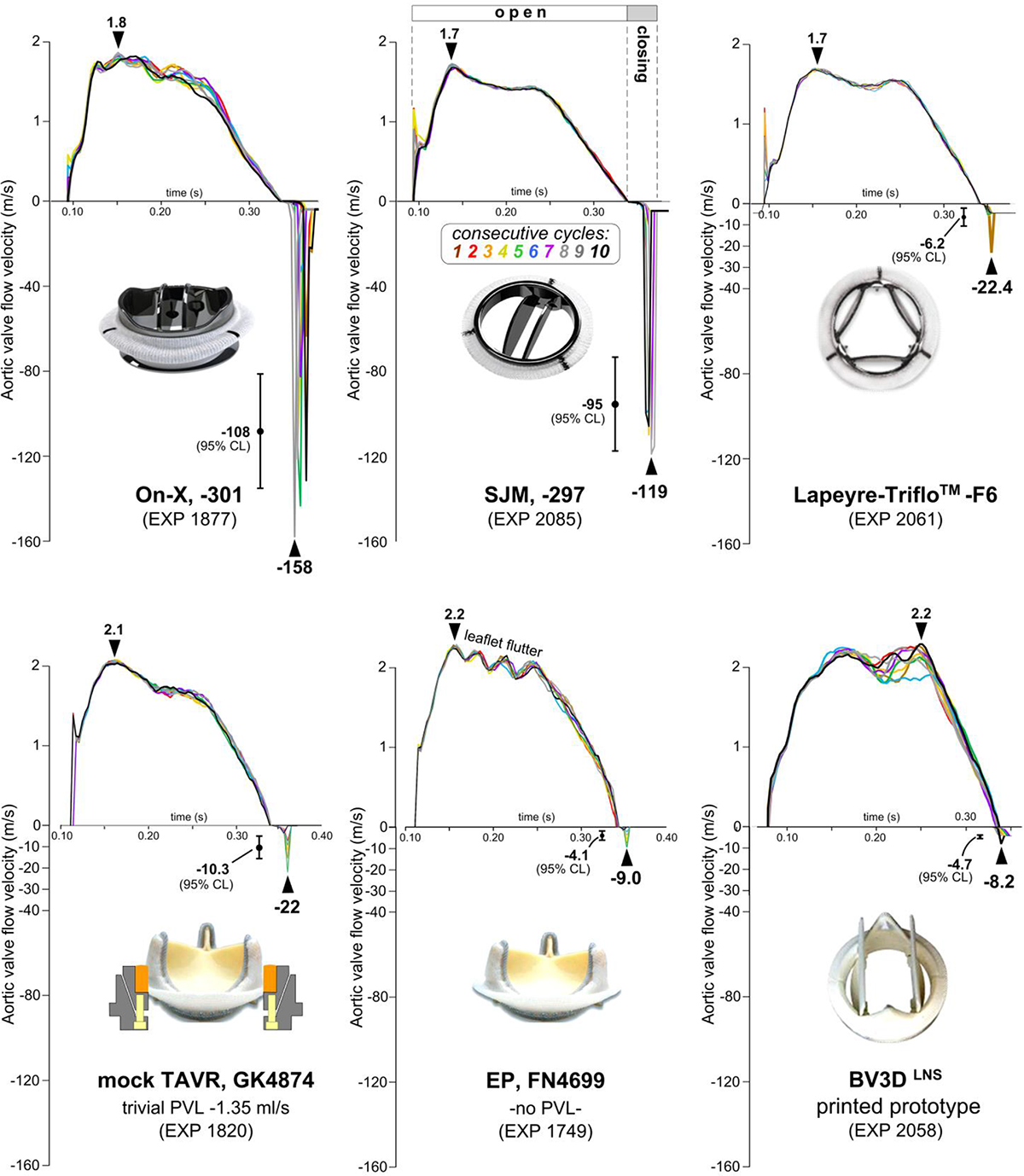
Laboratory data for various AORTIC valves of seat size 25mm. Flow velocities over 10- consecutive cycles with average, and 95% confidence limits identified (CL). Note, negative flow velocity transients during valve closure exceed the positive peak forward flow velocity. Results challenge the chronic emphasis (bias) towards forward flow open valve performance as the primary factor in valve-induced thrombogenic complications.

**FIGURE S4.**
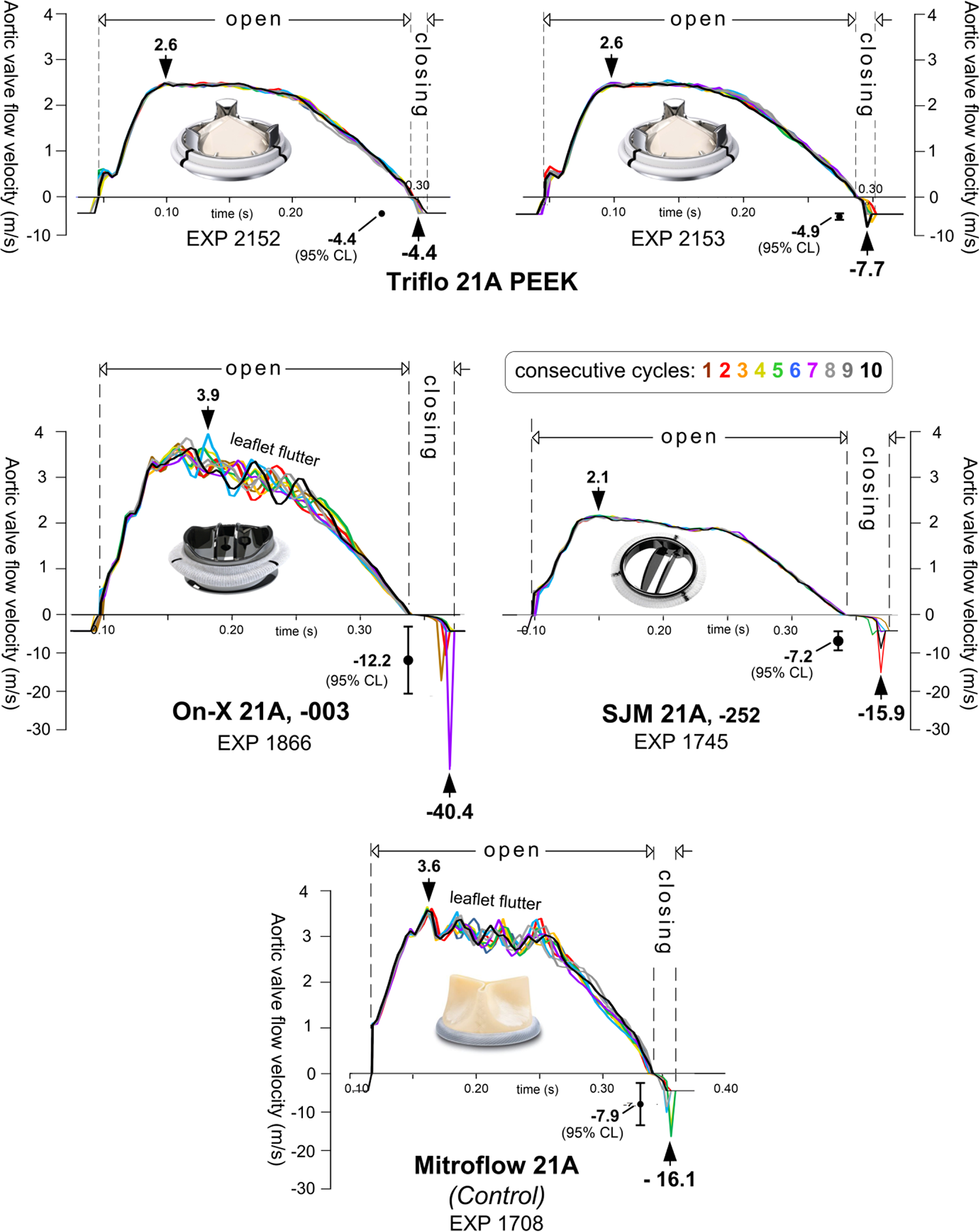
AORTIC valve flow velocity color coded profiles for 10 consecutive cycles are shown including average and 95% confidence limits (CL). Triflo PEEK valve profiles and data (top panel) for EXP 2152 and 2153 reveals favorable closing behavior and repeatability of experimental data under stable operating conditions. Note: negative transient flow velocities near valve closure exceed peak open flow phase velocities. Results challenge the conventional emphasis or bias that thrombogenic complications of valves is primarily induced by valve open phase performance.

## SPECIFICATIONS: (Scotten and Siegel 2014; 2015)

– Large area photodiode light sensitive area (10 square), Hamamatsu S1723-06
–Temporal resolution obtained from square-wave rise time ≈ 1.04 µs.

–Calibration reference areas *x*: 4.645, 3.577, 1.981, 1.005, 0.096 cm^2^
– Respective signal output values *y*: 3409, 2724, 1534, 772, and 75 A-D units
– EXCEL linear fit provides sensitivity equation:

*y* = 0.00134*x* A-D units
*x = y /* 0.00134 cm^2^
– Opto-electronics output ≈ 3.2 mv_rms_ (measured with light blocked by shutter)
– Spatial area resolution (based on measured baseline dark noise voltage):

–baseline dark noise = 3.2 mV_rms_ = 0.0032 V_rms_
–baseline dark noise (in A-D units) = 0.0032 V_rms_ × 204.8 A-D units/V_rms_ = 0.6554 A-D units Converting A-D units into area units = 0.6554 A-D units × 0.00134 cm^2^/A-D unit

=0.00088 cm^2^
=0.088 mm^2^

*For POA values below 0.088 mm², they are caused by various sources of error or uncertainty and are treated as insignificant noise rather than meaningful data*.

Maximum DC output from the opto-electronics is 10 Vdc (equivalent to 2048 A-D units) 1 A-D unit = 10 Vdc/2048 = 0.0049 Vdc

1V = 1/0.0049 A-D units = 204.8 A-D units

*y* (A-D units) = 0.00133 × noise 0.0032 V_rms_ = 0.000004277 A-D units

Equivalent area of noise signal *x* = 0.000004277 A-D units */* 0.00133 cm^2^ = **0.**0032 **cm^2^**

– Linear calibration (typical): y=0.00133*x*, with R^2^=0.9998.
–TML Telecentric lens with C-mount, Edmund Optics 56675, continuous magnification ≈0.16 ×
–Working distance ≈18 cm; Perspective error <0.3% (over depth of 15 mm). *Telecentric lenses have the unique property of maintaining a constant angle of view, even when the object’s distance changes. This is useful for precise measurements that avoid perspective error*.
– Variance of sensitivity over sensor area <6%;
–LED back light source (diffuse red): Phlox® SLLUB*****, 50×50mm x 8.5 mm, LED back light source

–Wavelength 625±15 nm.
–Luminance 5,780 cd/m^2^;
–Uniformity 99.24%;
*****PHLOX Corp., ZAC de l’Enfant, Aix-en-Provence, France

Summary of test results for size 21A aortic valves shown in Fig. S5.

1. EXPs 2152 and 2153, results are highly repeatable.
2. All valves have a very consistent behavior over 10 consecutive cycles especially for the Triflo and SJM valves with cycles essentially on top of each other.
3. On-X and Mitroflow valves have a prominent leaflet fluttering during the open phase.
4. Open times are about the same for all valves.
5. Closing times are equivalent between Triflo and Mitroflow but almost double for On-X and SJM.
6. Peak forward flow velocities are equivalent for the Triflo and SJM but are greater for On-X and Mitroflow.
7. Mean reverse-flow velocities are significantly lower for Triflo valve experiments and significantly higher for On-X with the SJM and Mitroflow in between.
8. Same observation for the peak reverse-flow velocities.

Figure. S5 shows Leonardo vs. animal aortic valve pressure gradients for Triflo, On-X/Cryolife, and Mitroflow 21A. The sheep (in vivo) data is reported by (Meuris et al. 2022).

## S7 Video

See MPEG 4, animated visualization of prototype BV3D valve structures and full range motion potential. https://youtu.be/5QmDPbfUvTMU-tube <u>link</u>

**Figure S5.**
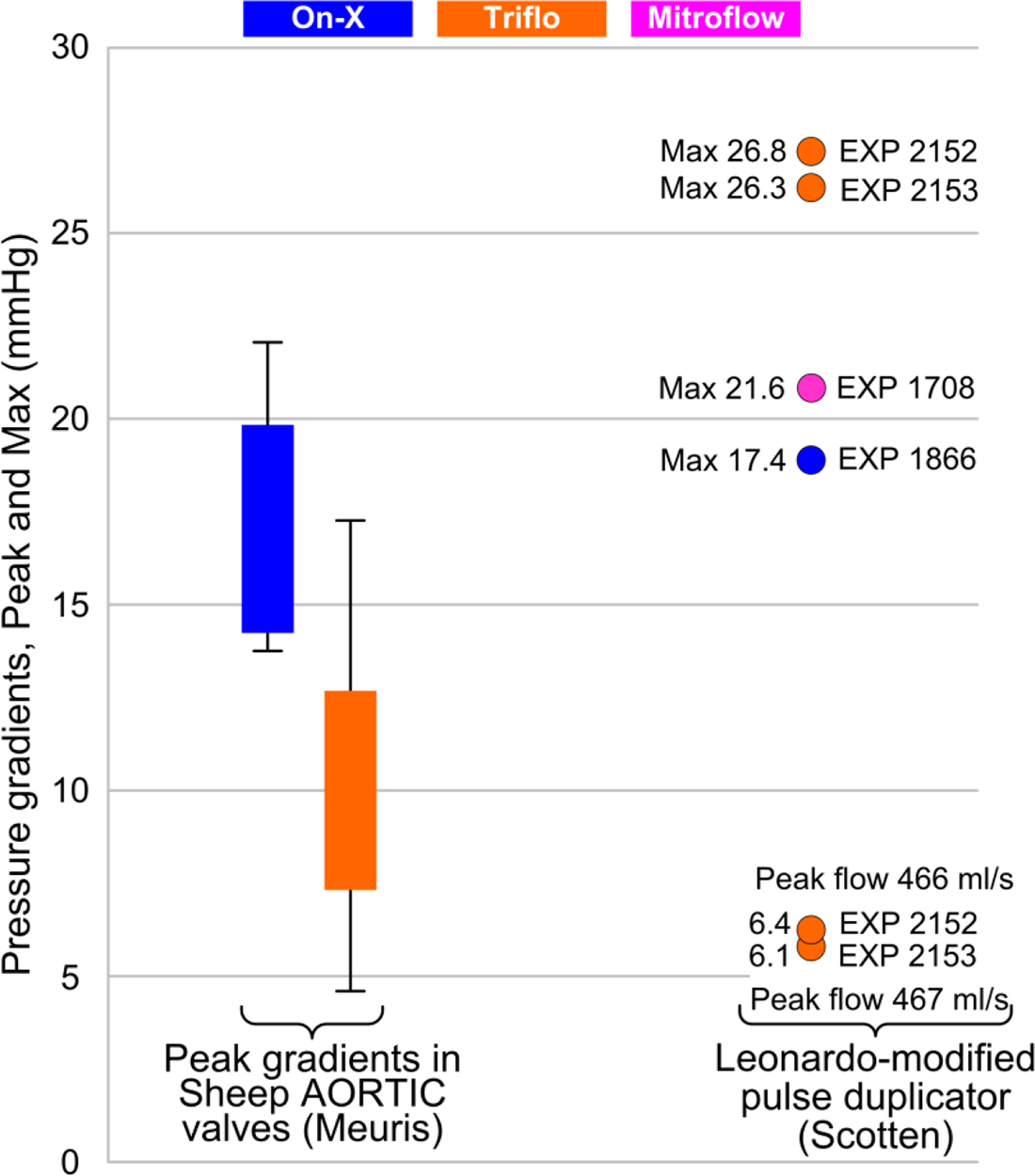
Clustered box plots show maximum *in vivo* pressure gradients. Circled data show *in vitro* pressure gradients (Max and @ Peak flows). Valves: On-X/Cryolife, Triflo PEEK and Mitroflow bioprosthetic. Valve seat size 21 mm Sheep aortic implants – (Meuris et al. 2022) Leonardo-modified pulse duplicator, (Scotten et al. 2022)

**Figure S6.**
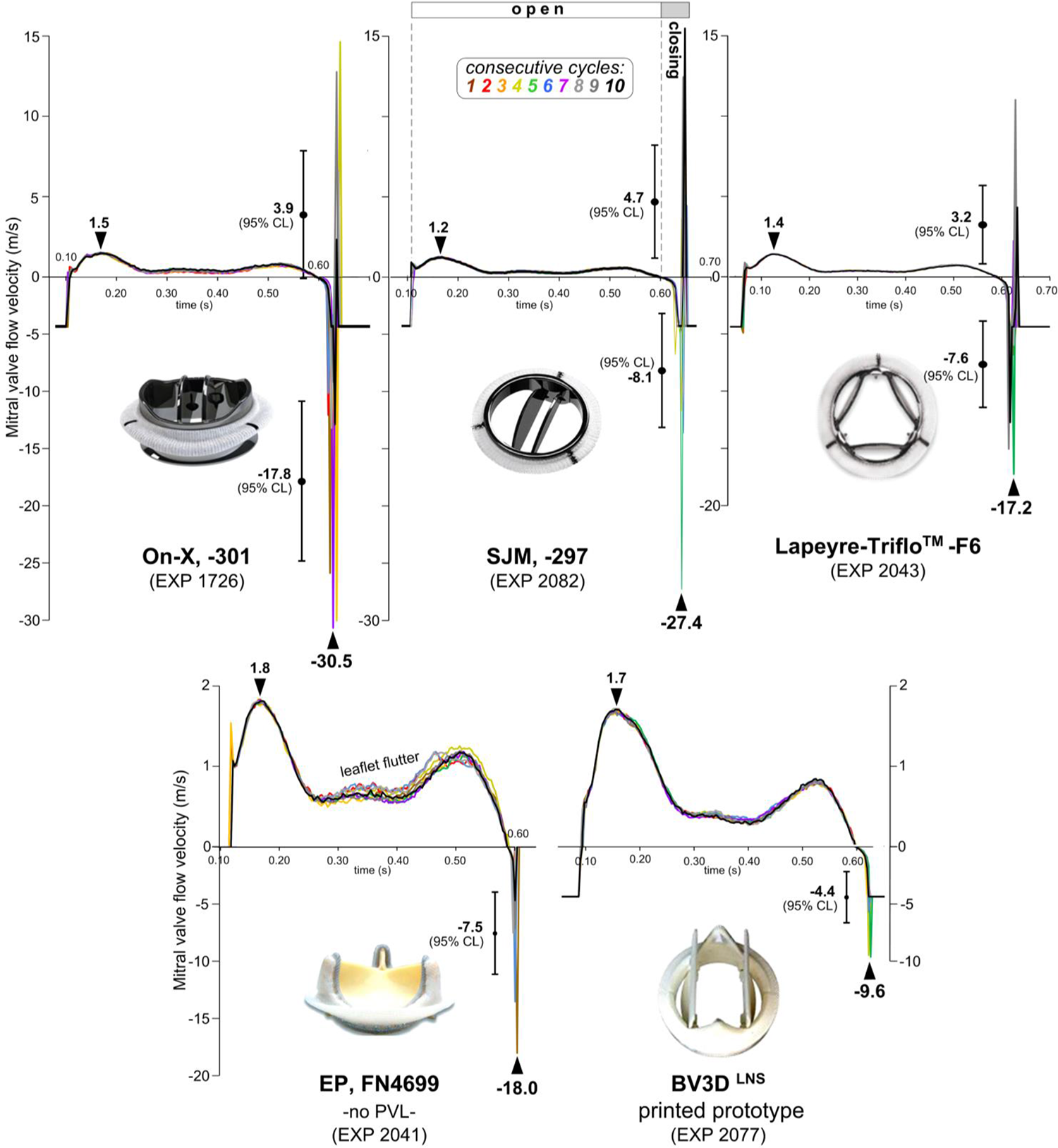
Laboratory data for MITRAL valve flow velocities over 10 consecutive cycles, average, and done to a 95% confidence limits (CL). Note, flow velocity transients during valve closure substantially exceed the maximum +’ve open valve flow velocities. Results challenge a persisting bias placed on open valve performance as the primary factor in valve-induced thrombogenic complications.

## S8 Reduced-order models

Figure 1 depicts Leonardo pulse duplicator system with reduced-order electrical models for the Viscoelastic Impedance Adapter (VIA) and outflow loading comprised of resistive and compliance elements used in the pulsed duplicator and assigned in computational fluid- structure simulations with saline used as the test fluid (Lee et al. 2020; Lee et al. 2021; Scotten et al. 2022).

Upstream VIA elements immersed in water:

> *C_VIA1_* = 0.0010 mL/mm Hg, 0.0275 mmHg mL^-1^, *C_VIA2_* = 0.1456 mL/mm Hg, 0.0347 mmHg mL^-1^

> *R*_VIA_ = 0.15 mm Hg/mL/s, 0.15mm Hg mL^-1^s**^2^**, and *R*_out_ = 0.0898mm Hg mL^-1^s. <u>Downstream elements:</u>

> *R*_c_ = 0.0218 mm Hg mL^-1^s, *R*_p_ = 1.31mm Hg mL^-1^s, and *C* = 1.27mm Hg mL^-1^. The test fluid is saline.

### Compliance and Resistance

A three-element (R–C–R) Windkessel model characterizes the upstream driving and downstream loading conditions specified in resistance and compliance units:

> –Upstream:

> *C*_VIA1_ = 0.0275 mm Hg mL^-1^, *C*_VIA2_ = 0.0347 mm Hg mL^-1^,

> *R*_VIA_ = 0.15mm Hg mL^-1^s, and *R*_out_ = 0.0898mm Hg mL^-1^s

> –Downstream:

> *R*_c_ = 0.0218 mm Hg mL^-1^s, *R*_p_ = 1.31mm Hg mL^-1^s, and *C* = 1.27mm Hg mL^-1^

Mathematically, compliance is defined as the ratio of change in air volume to change in air pressure and is considered a bulk modulus of elasticity or young’s modulus of air (E):

> C =1/E (in SI units of dyne/cm)

Compliance is modeled by four enclosed air volumes:

> *C_VIA1_* = 120 ml (0.0010 mL/mm Hg)

> *C_VIA2_* = 50 ml (0.1456 mL/mm Hg)

> (*C_root_*) = 640 mL Aortic root

> *C_per_* = 615 ml Systemic arterial compliance

Compressibility of air is calculated by a bulk modulus (K):

> K=1333.2 × VΔP/ ΔV ≈ 140 kPa⁻¹ for standard air conditions Where:

> Given that K ≈ 140 kPa⁻¹ for standard air conditions, Check that calculated K is to this approximate value. V is initial contained air volume (cm^3^)

> ΔP is the change in contained air pressure (mm Hg)

> ΔV is the resulting change in contained air volume (cm^3^) Conversion factor: 1 mmHg =1333.2 dynes/cm^2^

The bulk pump source compliances simulated in VIA contain two air volumes *C_VIA1_* and *C_VIA2_*. These volumes are adjustable and cover the physiological range. The nominal air volume settings used for valve testing are: *C_VIA1_* = 120 ml (0.0010 mL/mm Hg); *C_VIA2_* = 50 ml (0.1456 mL/mm Hg); aortic root (*C_root_*) = 640 mL and systemic arterial compliance *C_per_* = 615 mL.

Compliance is defined as the ratio of volume change to pressure change as follows:

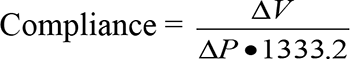

Where:

Δ*V* = change in contained air volume in ml

Δ*P* = change in pressure (mm Hg) caused by volume change Δ*V*

Δ*P* = *P*_2_ - *P*_1_

*P*_1_ = initial static pressure in mmHg

*P*_2_ = final static pressure in mmHg

Conversion factor: 1 mmHg = 1333.2 Dynes/cm^2^

Air volume_max_ values found experimentally that simulate left ventricle, aortic root, and systemic arterial compliance and calibrated parameters for the reduced-order models using saline as a test fluid (Lee et al. 2020; Lee et al. 2021; Scotten et al. 2022) facilitated realistic pressure and flow wave forms while under pulsatile flow conditions were:

- R_VIA_ = 0.15 mm Hg/mL/s.
- Aortic root *C_VIA1_* = 120 ml = 0.1456 mL/mm Hg
- Output compliance *C_VIA2_* = 50 ml
- Aortic root *C_root_* = 640 ml= 0.0010 mL/mm Hg
- Left ventricle source compliance air volume *C_VIA2_* = 640 ml
- Output compliance air volume = 50 ml
- Peripheral systemic *C_per_* = 615 ml

### Resistance

Resistance to flow causes frictional loss of energy and flow container chamber radius is the dominant determinant of resistance. In Figure 1, R_VIA_ and R_per_ offer flow resistance. R_VIA_ consists of a micro-porous water filter section which offers a low fixed resistance to flow (200 c.g.s. units). The peripheral resistance R_per_ offers alterable resistance allowing for operator adjustment of end diastole aortic pressure.

### *S9

A mining tradition dating back to 1911 used canaries in coal mines to detect carbon monoxide and other toxic gases before they hurt humans. In 1906, Dr. John Scott Haldane proceeded to study asphyxia in coal miners in coal miners and proposed that *“miners carry small animals like mice or canaries to detect levels of gas in their working environment”* This practice was in vogue until 1986. (Sekhar KC, Rao SC. John Scott Haldane: The father of oxygen therapy. Indian J Anaesth 2014;58:350-2. <u>Link</u>

